# IntegrateALL: an end-to-end RNA-seq analysis pipeline for multilevel data extraction and interpretable subtype classification in B-precursor ALL

**DOI:** 10.1101/2025.09.25.673987

**Authors:** Nadine Wolgast, Thomas Beder, Mayukh Mondal, Wencke Walter, Stephan Hutter, Sonja Bendig, Jan Kässens, Björn-Thore Hansen, Katharina Iben, Sebastian Wolf, Anjali Cremer, Malwine Barz, Martin Neumann, Nicola Gökbuget, Claudia Haferlach, Monika Brüggemann, Claudia D Baldus, Alina M. Hartmann, Lorenz Bastian

## Abstract

Transcriptome sequencing (RNA-seq) is emerging as a diagnostic standard for B-cell precursor acute lymphoblastic leukemia (B-ALL). Expression-based classifiers reach ∼95% accuracy, but reproducible end-to-end solutions that also integrate transcript-derived genomic drivers and quantitative virtual karyotyping are lacking. We developed IntegrateALL, a Snakemake pipeline that standardizes RNA-seq analysis from FASTQ to rule-based subtype assignment across 26 WHO-HAEM5/ICC entities by integrating expression-based subtype prediction, gene fusion- / hotspot SNV calling and virtual karyotyping. We introduce KaryALL, a machine-learning classifier that uses normalized expression and minor-allele-frequency features (RNASeqCNV) to distinguish near haploid, hypodiploid and high hyperdiploid B-ALL and chromosome-21 gains/iAMP21 (accuracy: 0.98 / F1-score: 0.96 on 615 independent test samples). SNP-array concordance supported RNA-based karyotyping. Applied to 774 unselected B-ALL cases, IntegrateALL yielded unambiguous subtype assignments in 81.5%, based on concordance of gene expression class with a defining driver (75.3% of all cases) or, in selected cases, high-confidence expression-based classification alone (6.2%); the remainder (18.5%) were flagged for manual curation. Independent validation (3 cohorts; n=436, including pediatric cases) reproduced these distributions. Across all patients (n=1,210), 2.6% harbored two subtype defining drivers, including hyperdiploidy in fusion-driven subtypes where it was not expected or subtype-defining SNVs (e.g., *PAX5* P80R / *IKZF1* N159Y) co-occurring with *BCR::ABL1*-positive/-like, *KMT2A*- or *DUX4*-fusions. In most dual-driver cases, one subtype gene expression signature predominated, indicating a hierarchy of oncogenic control and the value of systematic driver screening alongside expression-based calls. IntegrateALL provides an adaptable fully reproducible workflow for molecular B-ALL characterization by systematically integrating genomic drivers and downstream gene regulation.

## Introduction

Diagnostic accuracy in hematology relies on systematic and reproducible workflows. The current World Health Organization Classification of Haematolymphoid Tumours (WHO-HAEM5)^1^ and the International Consensus Classification (ICC)^2^ of Myeloid Neoplasms and Acute Leukemias recognize 12 and 22 molecular subtypes of B precursor acute lymphoblastic leukemia (B-ALL), respectively, as distinct diagnostic entities. Additionally, the ICC defines five provisional subtypes. Diagnostic definitions are based on recurrent genomic aberrations such as specific aneuploidy patterns, gene fusions, single-nucleotide variants, and complementary gene expression signatures. These expression signatures typically reflect either the transcriptional regulation driven by subtype-specific genetic alterations or integrated regulatory networks downstream of multiple driver aberrations (e.g., *BCR::ABL1*-like ALL, *PAX5*alt ALL). Further subclusters with these subtypes reflect functional and clinical heterogeneity (e.g., multilineage vs. lymphoid *BCR::ABL1*-positive ALL and JAK/STAT vs. ABL class driven *BCR::ABL1*-like ALL).

Existing subtype definitions, however, have mostly been characterized in specifically selected cohorts or age-specific contexts. Age-overriding systematic catalogues of driver aberrations and corresponding gene expression signatures remain lacking.

Transcriptome analysis by RNA-sequencing (RNA-Seq) allows comprehensive profiling of gene expression together with inference of underlying genomic drivers. A variety of bioinformatic tools have been established for tasks such as gene expression count quantification^3^, gene fusion calling^4,5^, and identification of expressed single-nucleotide variants^6,7^. We and others have developed approaches for systematic gene expression-based subtype classification^8–13^. Accurate subtype classification requires integration of this multi-level data, accounting also for the rare possibility of multiple drivers in a single sample. Some existing classifiers^11–13^ already use outputs from multiple tools for interactive visualization and classification. However, comprehensive and transparent end-to-end workflows from raw FASTQ-files to rule-based classifications according to current diagnostic definitions are lacking, making subtype allocation resource-intense and prone to errors during manual curation.

Certain molecular subtypes, such as Near Haploid, Low Hypodiploid, and hyperdiploid ALL are defined by patterns of non-random chromosomal gains and losses. They represent a particular challenge for RNA-based classification due to the absence of a single defining driver lesion and highly similar gene expression patterns between Near Haploid and hyperdiploid ALL^8^. RNASeqCNV^14^ addresses this challenge by reconstructing virtual karyotypes from normalized gene expression count data and expressed variant allele frequencies. However, the definition of these aneuploidy-based subtypes currently relies on frequency distributions rather than clear quantitative criteria, and methods to objectively assess a sample’s karyotype against these distributions are not available. Thus, karyotype interpretation remains to some extent subjective, limiting reproducibility.

To address these gaps and enable systematic and reproducible subtype allocation, we developed IntegrateALL, a free open-source Snakemake^15^ pipeline. IntegrateALL provides a standardized workflow from raw RNA-Seq FASTQ files through quality assessment, gene expression quantification, identification of gene fusions and hotspot SNVs, and reconstruction of virtual karyotypes. Classification according to current diagnostic criteria is implemented via a novel, rule-based system informed by own^8,16,17^ and published data^18–20^ on driver aberrations and corresponding gene regulation. For an automated quantitative analysis of B-ALL karyotype patterns, we developed and implemented KaryALL, the first machine learning classifier for RNA-Seq based virtual karyotypes. Easy accessibility of all data layers in an interactive HTML output ensures transparency for the classification process and offers a user-friendly environment for manual curation of complex cases which do not fit standard diagnostic definitions. Validation on 1,210 B-ALL samples from four cohorts across age groups (Supplementary Table S1) confirmed the scalability and robustness of our pipeline. Its open design ensures adaptability to evolving diagnostic criteria and systematic analytical workflows in B-ALL.

## Materials and Methods

To establish IntegrateALL as a systematic B-ALL RNA-Seq analysis workflow, we used Snakemake^15^, a Python-based workflow management system, to integrate raw data quality check (FASTQC^21^ and MULTIQC^22^), read alignment^3^ to GRCh38.83, gene fusion calling (ARRIBA^5^ and FusionCatcher^4^) and raw single-nucleotide variant calling (GATK, Genome Analysis Toolkit)^7^. High-confidence hotspot SNV were filtered with pysamstats^6^ against a curated list of B-ALL relevant codon changes (*ZEB2*, *KRAS*, *NRAS*, *FLT3*, *PAX5*, and *IKZF1*; Supplementary Table S2). RNASeqCNV^14^ was used to infer virtual karyotypes from normalized counts and expressed variant allelic frequencies derived from GATK calls^7^. The GATK workflow, executed with Snakemake wrappers, adheres to best practices for RNA-seq short variant discovery, including read group addition, duplicate marking, exon segmentation, base recalibration, and variant filtration. For a systematic classification of aneuploid B-ALL subtypes, we developed KaryALL, which uses chromosome-level expression features and minor-allele-frequency distributions from RNA-Seq CNV outputs. Models were built in Python using scikit-learn^23^ (data preprocessing, RandomForestClassifier, KNeighborsClassifier) imblearn^24^ for class balancing via SMOTE^25^, and XGBClassifier from xgboost package^26^. For systematic gene expression-based subtype allocation we included ALLCatchR^8^, our machine learning based classifier which provides automated allocation to 21 B-ALL subtypes and reports proximity to normal lymphopoiesis, inferred immunophenotype, sex, and blast proportion. Outputs from individual tools were integrated using a newly established rule set reflecting genomic / transcriptomic definitions consistent with current diagnostic definitions.^2^/ ^1^

We tested IntegrateALL on n=774 B-ALL first diagnosis bone marrow / peripheral blood samples from adult patients extending our previously published cohort of patients treated on protocols of the German Multicenter Study Group for Adult ALL (GMALL)^17^. Patients consented on the scientific use of diagnostic leftover materials (Kiel University Ethics committee vote D416/21). RNA-Seq was performed as described previously^16,17^. Briefly, libraries were prepped using TruSeq or Illumina Stranded mRNA Prep (Illumina, San Diego, USA), and sequenced aiming for 30 Mio paired-end reads (75 – 150 bp) on a NovaSeq or NextSeq sequencing system (Illumina, San Diego, USA). For orthogonal genomic karyotyping, DNA from diagnostic materials was used for SNParray profiling (Infinium Global Screening Array-24 v3.0, Illumina, San Diego, USA). Array raw data were processed in Genome Studio (Illumina, San Diego; USA, v2.0.5) to derive allelic depth and variants. For further validation of KaryALL and IntegrateALL, we used real-world diagnostic RNA-Seq data with a genomically validated ground truth^27^, and published RNA-Seq data^28–30^ with a specific focus on aneuploid subtypes which were identified by agreement between ALLCatchR and ALLSorts^31^. IntegrateALL is available as a free open-source pipeline under the MIT License (https://github.com/NadineWolgast/IntegrateALL).

## Results

### Development of a computational framework for systematic subtype allocation in B-ALL

RNA-Seq data contains multiple data levels, which are informative for molecular subtype classification. In B-ALL, subtypes are defined by consistency between gene expression signatures and genomic drivers inferred from RNA-Seq profiles. To systematically extract these features in an accessible manner, we developed IntegrateALL, a Snakemake pipeline (Figure 1), which starts from RNA-Seq FASTQ files to perform the following analyses: read quality assessment, gene expression-based subtype allocation, gene fusion calling, identification of hotspot single nucleotide variants and analysis of virtual karyotypes. Data are made accessible via an interactive HTML report and are used for integrative subtype classification using a rule set based on current diagnostic definitions^1,2^.

**Figure 1:**
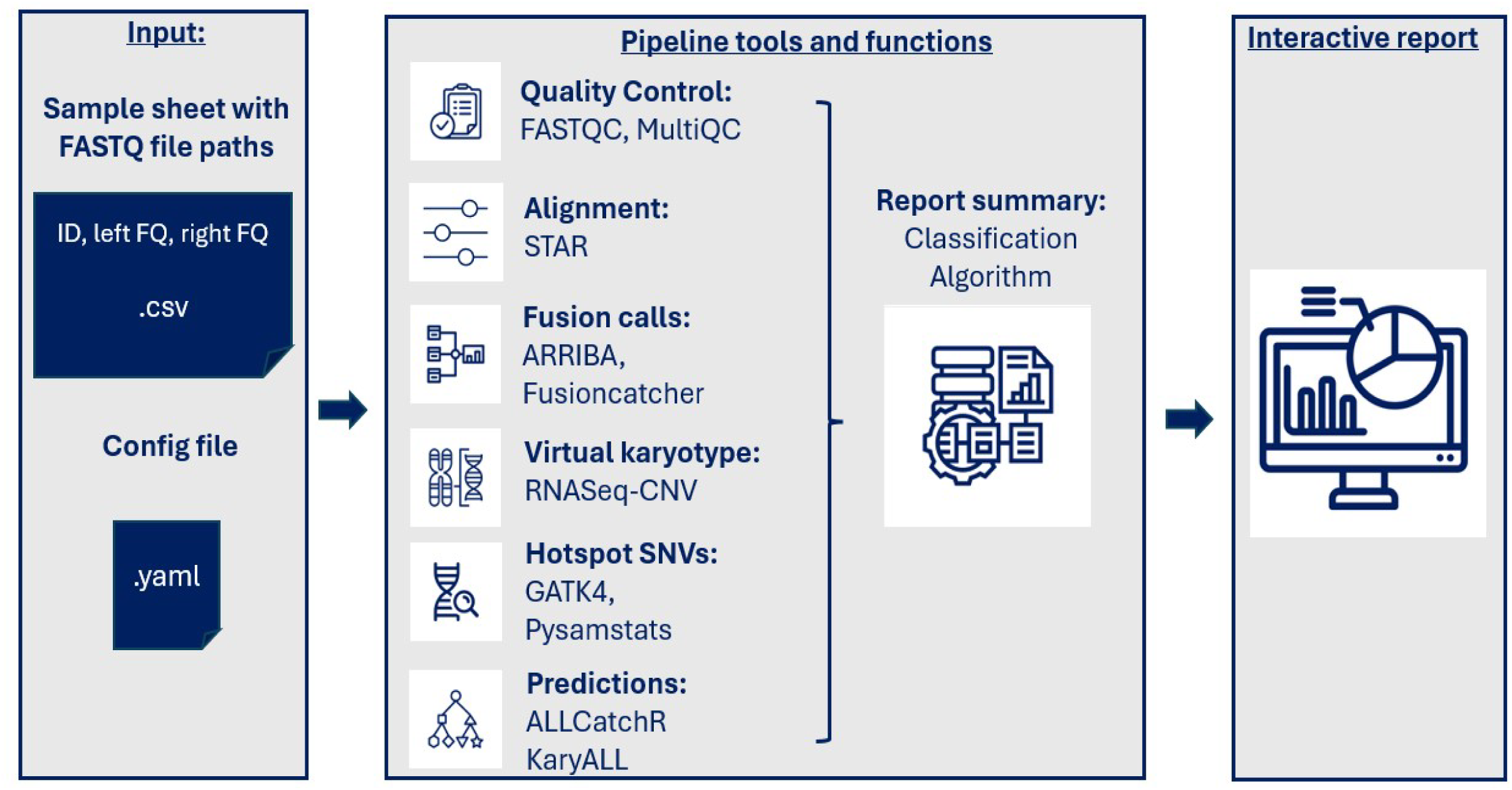
Schematic overview of the IntegrateALL pipeline. The workflow starts with an input sample sheet and raw RNA-seq FASTQ files. Subsequent steps integrate quality control (FASTQC^21^, MULTIQC^22^), read alignment (STAR^3^), gene fusion detection (ARRIBA^5^, FusionCatcher^4^), SNV calling (GATK^7^, pysamstats^6^), virtual karyotyping (KaryALL classification based on RNASeqCNV^14^ outputs), and gene expression–based subtype classification (ALLCatchR^8^). A configuration file enables resource optimization and parameter customization. An interactive summary output integrates results from all tools for manual review and displays the final classification (WHO-HAEM5/ICC).

### KaryALL: a Machine Learning Classifier for aneuploid B-ALL subtypes

Non-random chromosomal gains and losses define Near Haploid (23-29 chromosomes), Low Hypodiploid (33-39 chromosomes) and High hyperdiploid (52-67 chromosomes)^32^ ALL. Near Haploid and High hyperploid subtypes share highly similar gene expression patterns, representing a challenge to gene expression-based subtype classification. Intrachromosomal amplification of chromosome 21 (iAMP21) is a structural aberration sufficiently large to be detectable by virtual karyotyping. It defines a molecular B-ALL subtype with a gene expression signature highly similar to *BCR::ABL1*-like ALL. To facilitate systematic classification of these four molecular subtypes, we used RNASeqCNV^14^ outputs of chromosome-specific gene expression counts and minor allele frequency distributions (Figure 2A) to train and validate a machine learning classifier. For training, we used an aggregated RNA-Seq data set of 395 samples from 7 cohorts classified according to published ground truth and ALLCatchR^8^ / ALLSorts^31^ predictions as: Near Haploid (n=6), Low Hypodiploid (n=28), hyperdiploid (n=75), iAMP21 (n=7) or non-aneuploid (‘other’; n=279; Supplementary Figure 1). For comparison with genomic karyotypes, 33 samples (n=11 hyperdiploid, n=11 low hypodiploid and n=11 diploid karyotypes) underwent SNParray profiling. RNASeqCNV calls were concordant in n=719/759 chromosomes analyzed (Figure 2B; p=5.00E-4), supporting the applicability of RNASeqCNV for virtual karyotyping.

**Figure 2:**
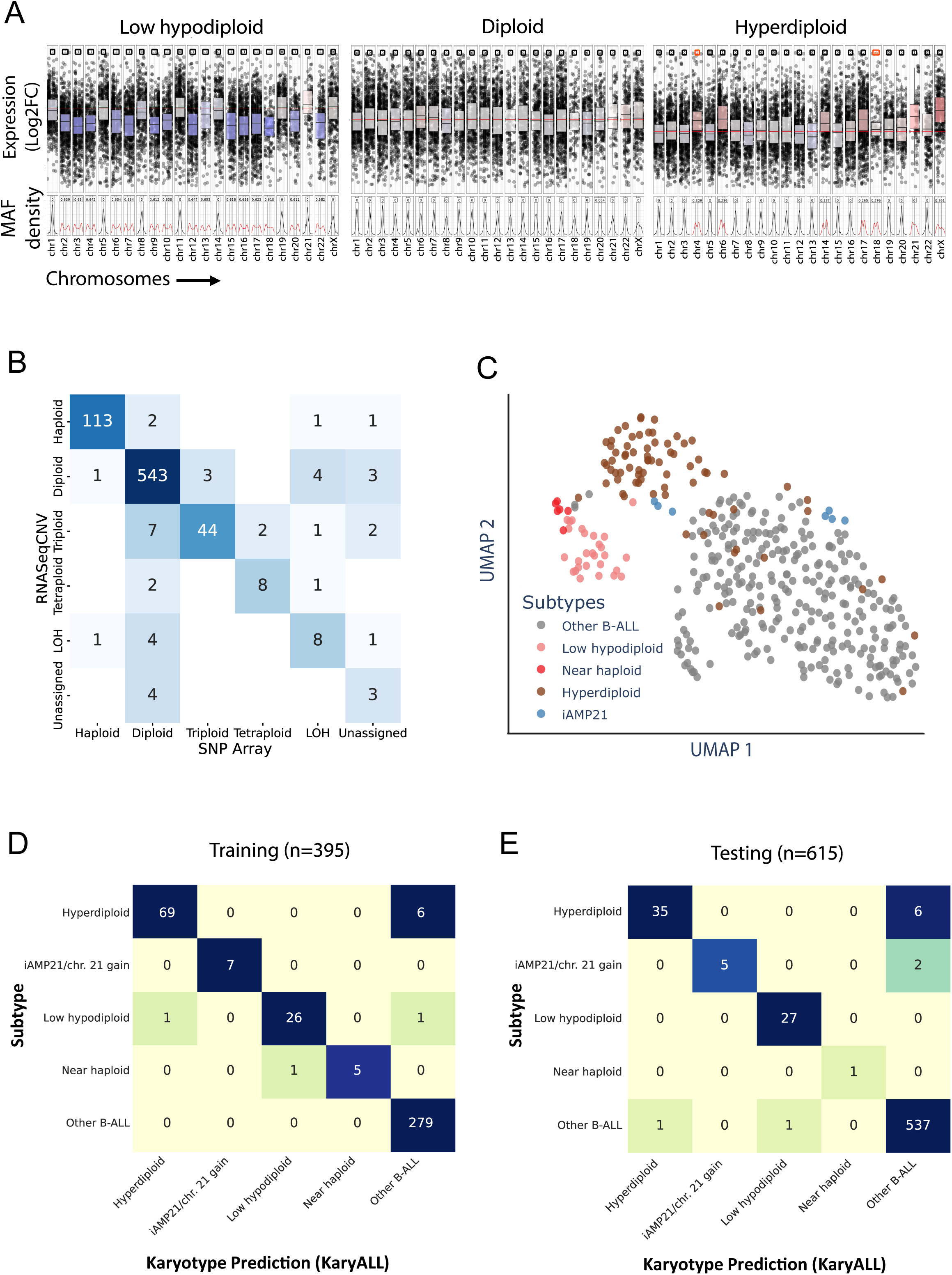
Training and validation of the KaryALL virtual karyotype classifier. (A) RNASeqCNV provides virtual karyotypes from RNA-seq data leveraging normalized counts and variant allele frequency (MAF) densities. Plots illustrate representative low hypodiploid, diploid, and High hyperdiploid cases. (B) Ploidy states and loss of heterozygosity (LOH) of individual chromosomes (n=33 samples) were determined by manual curation of RNASeqCNV outputs and compared to SNParray outputs. (C) B-ALL samples (n=395) with available SNParray and RNA-seq data were analyzed by Uniform Manifold Approximation and Projection for Dimension Reduction (UMAP) using RNASeqCNV normalized expression and MAF density data. Colors indicate molecular subtypes according to genomic ground truth and manual curation of RNASeqCNV / SNParray outputs. This data set was used for training the machine learning classifier KaryALL to predict B-ALL karyotypes based on RNASeqCNV outputs. The corresponding comparison of RNASeqCNV and SNParray profiles and details on the KaryALL design are shown in Supplementary Figures S1-3. (D) Confusion matrix of KaryALL classification accuracy on the training data set (accuracy: 0.98 / F1-score: 0.96). (E) Confusion matrix of KaryALL classification accuracy on test data from three independent cohorts (accuracy: 0.98 / F1-score: 0.98).

Unsupervised UMAP analysis separated most subtypes (Figure 2C) with the least distinct clustering for iAMP21. To improve iAMP21 performance, we defined 29 additional chromosome 21-specific count features and minor allele frequency values for incorporation into training. Analysis of rare non-iAMP21 cases with isolated chromosome 21 amplification and otherwise diploid karyotypes indicated that these could not be distinguished in RNASeqCNV outputs from iAMP21. Therefore, we considered these as a combined class,iAMP21/chr21-amplification‘. SMOTE upsampling increased the representation of iAMP21 and Near Haploid subtypes to approximate Low hyperdiploid ALL frequency. We established an ensemble soft-voting classifier from the three best performing machine learning models (Random Forest, K-Nearest Neighbors (KNN), and XGBoost; Supplementary Figure 2). In leave-one-out cross-validation, individual models achieved accuracies of 0.95-0.97 and F1 scores of 0.82-0.93 (Supplementary Figure 3). Hyperparameter tuning was conducted to optimize performance, particularly for underrepresented subtypes such as iAMP21 and Near Haploid cases. The ensemble classifier achieved an accuracy of 0.98 and a F1 score of 0.96 using leave-one-out cross validation (Figure 2D). The average sensitivity and specificity for detection of aneuploid karyotypes or iAMP21/chr 21 amplification were 0.99 and 0.92, respectively. The lowest performance was observed for near-haploid cases (sensitivity 1.00, specificity 0.83; n = 6), while the highest accuracy was achieved for iAMP21 chr 21 amplification (sensitivity and specificity both 1.00; n = 7). Robustness was confirmed in two independent cohorts (n=105 Munich Leukemia Laboratory; n=206 pediatric B-ALL samples^28^) and our own cohort (n=304), totaling 615 samples (Figure 2E). The overall weighted F1-Score was 0.98 and the accuracy was 0.98 as well, demonstrating the model’s robustness and its applicability to broader datasets.

To our knowledge, KaryALL is the first systematic classifier for virtual B-ALL karyotypes and has therefore been implemented to complement gene expression-based subtype allocation with the corresponding genomic driver profile.

### Classification for 26 molecular B-ALL molecular subtype definitions

Current hematologic disease classifications define 12^1^ and 22^2^ B-ALL molecular subtypes as well as 5 provisional entities. We formalized B-ALL subtype allocation within our pipeline by establishing a rule-set for systematic integration of (i) our gene expression-based reference for 21 B-ALL subtypes^8,33^, (ii) an expanded catalogue of 165 driver gene fusions driver candidates based on own^16^ and published data sets^16,19,20^, (iii) KaryALL karyotype classifications and (iv) subtype-defining single-nucleotide variants (e.g., *PAX5* p.P80R or *IKZF1* p.N159Y; Figure 3, Supplementary Figure 4, Supplementary Table S3).

**Figure 3:**
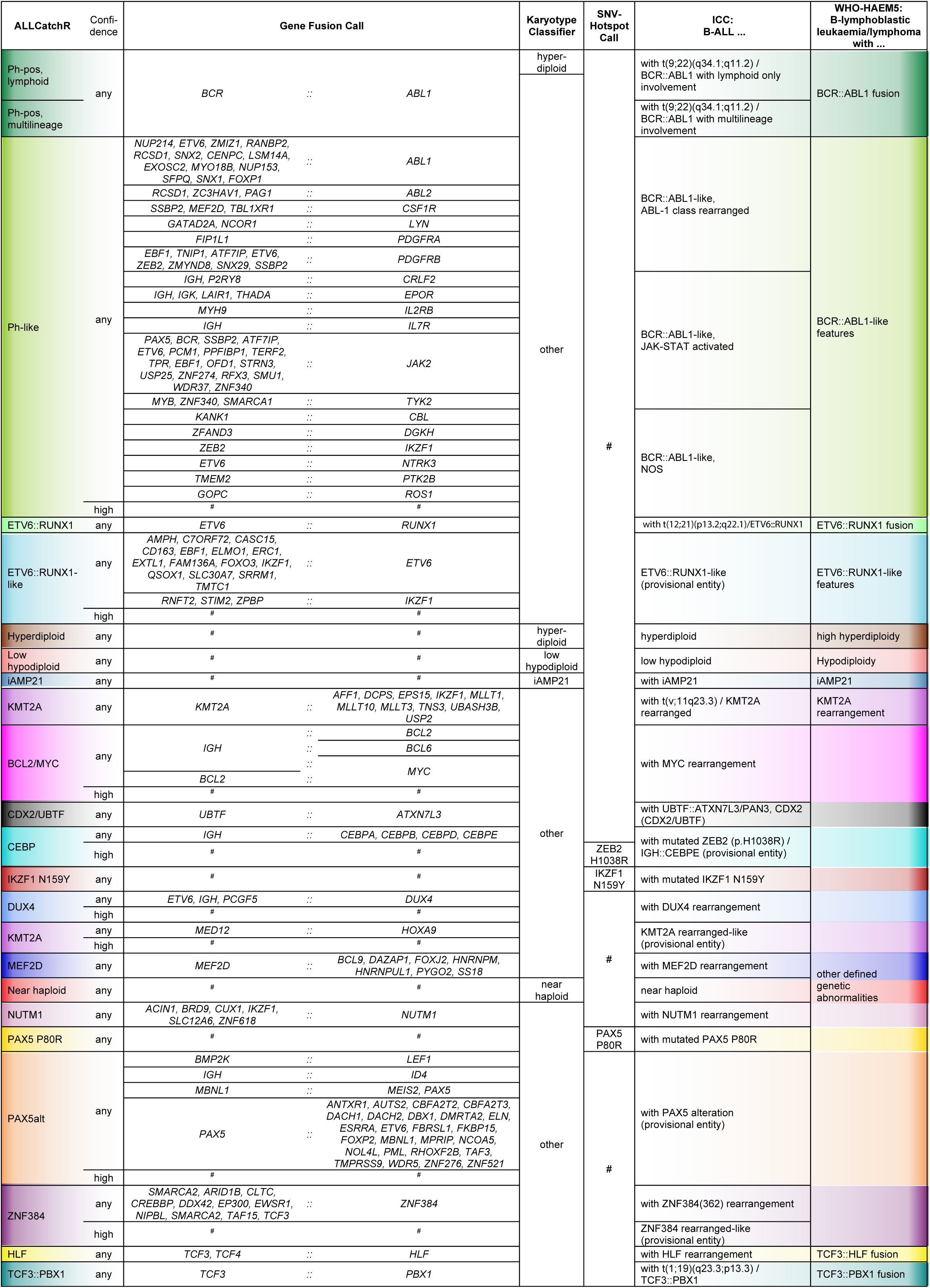
B-ALL subtype definitions. Molecular B-ALL subtype definitions used for subtype classification in IntegrateALL. This overview represents an integration of gene expression-based subtype (ALLCatchR), gene fusion calls, virtual karyotype classification (KaryALL), and recurrent hotspot single nucleotide variants. The entire rule set, and the classification logic are presented in Supplementary Table S3 and Supplementary Figure S4. Unambiguous matches to this rule set are automatically classified according to current classifications (ICC / WHO-HAEM5). ‘#’ indicates classes where subtype defining SNVs are absent.

For an unambiguous subtype allocation, we require concordance between a subtype-defining genomic driver and the corresponding ALLCatchR gene expression subtype and absence of any other subtype-defining driver aberration (Figure 3). For such cases any ALLCatchR confidence level was accepted. Because original definitions included gene-expression-only allocations for certain subtypes (e.g., *BCR::ABL1*-like, NOS^31^ and *PAX5*alt^19^) and because gene fusions involving the IGH locus pose a specific challenge for identification by RNA-Seq, we allowed high confidence expression-only calls for IGH-fusion driven subtypes (i.e. *DUX4*, *CEBP*) or *BCR::ABL1*-like and *PAX5*alt provided no alternative driver was present. All samples fitting these definitions were automatically classified according to n=10/11 WHO-HAEM5 and n=26/27 ICC molecular subtypes; *IGH::IL3* rearranged B-ALL was not represented due to its rarity and lack of gene expression data. The remaining cases were flagged for manual curation. An overall open rule set design permits flexible adaptation to incorporate novel subtype definitions.

### IntegrateALL yields concordant expression and driver/karyotype calls in 80% of B-ALL RNA-Seq profiles

We applied IntegrateALL to n=774 unselected adult B-ALL cases, comprising samples from our previously published dataset^16^ and additional cases from ongoing RNA-Seq-based routine diagnostics (Figure 4). Of all patients, 43.9% were female and 56.1% male, with 14.3% and 85.7% pro-B versus pre-B/common inferred ALL immunophenotype and with an overall median blast proportion of 72.8% according to ALLCatchR predictions. ALLCatchR provided high-confidence gene expression subtypes in n=560/774 cases (72.3%) and candidate subtypes in n=181/774 cases (23.4%); n=33/774 patients (4.3%) remained unclassified by expression alone. KaryALL identified Near Haploid (n=2), Low Hypodiploid (n=42), High hyperdiploid (n=46) and iAMP21 (n=1) karyotypes, confirmed by manual curation of RNASeqCNV outputs. Subtype-defining driver fusions were detected in 476 samples (54 unique drivers; Supplementary Table S4, Supplementary Figure 5). Recurrent cooperating hotspot events included *NRAS* (n=121/774; 15.6%), *KRAS* (n=67/774; 8.7%) and *FLT3* hotspot SNVs (n=40/774; 5.2%).

**Figure 4:**
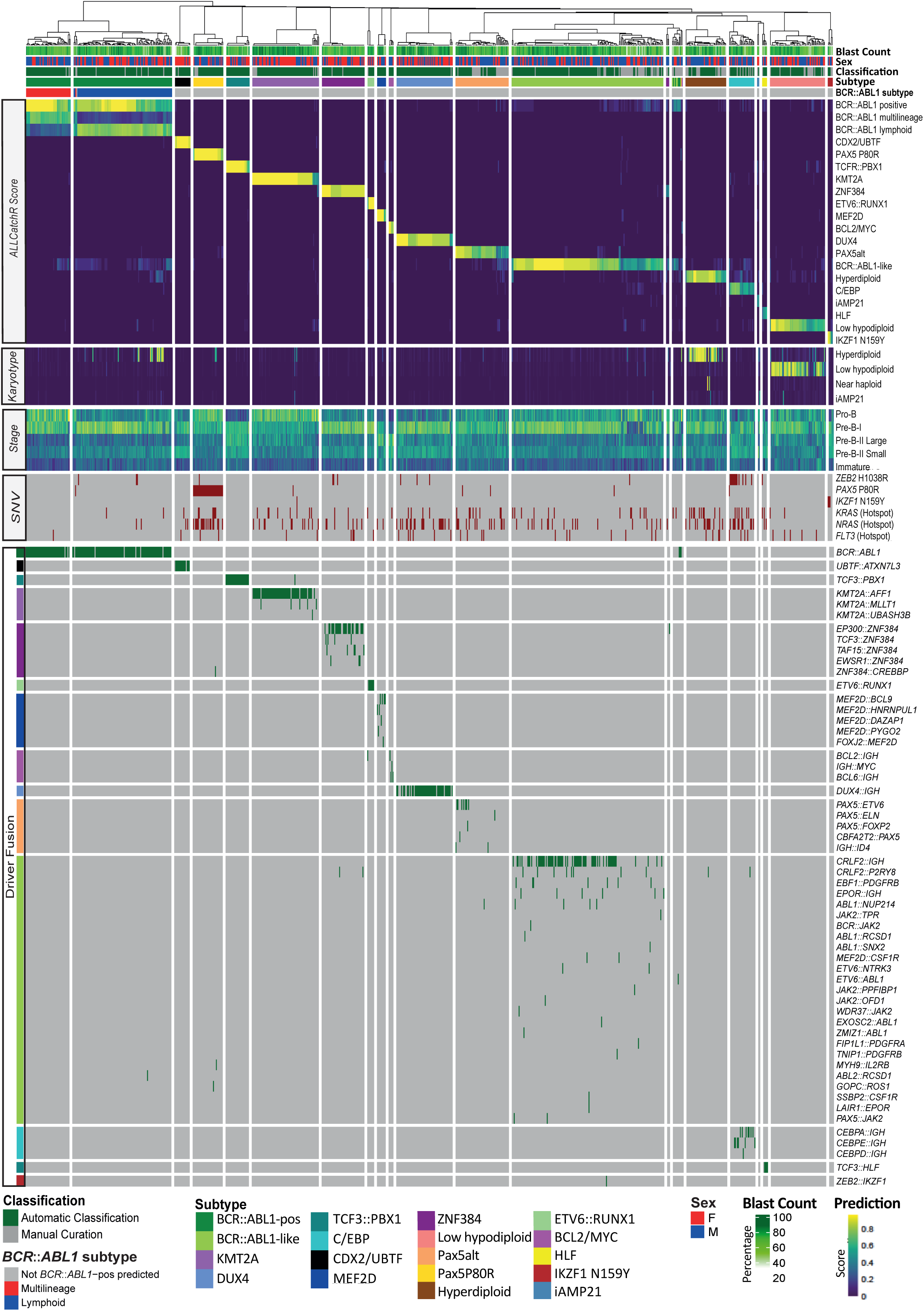
IntegrateALL classification of unselected adult B-ALL RNA-seq samples. A total of 774 B-ALL samples from own sequencing were processed through IntegrateALL, yielding an overview of gene expression based subtype prediction (ALLCatchR), virtual karyotype classification (KaryALL), proximity to normal B cell development (ALLCatchR), hotspot single nucleotide variants (GATK workflow, Pysamstats) and driver fusion calls (FusionCatcher, ARRIBA) together with patient’s sex and blast count predictions (ALLCatchR) and the IntegrateALL annotation for automated classification vs. manual curation.

Application of our defined rule set (Figure 3) to our B-ALL cohort provided automated subtype allocations in n=631/774 (81.5%) of cases (Figure 5A) based on concordance between gene expression subtype and gene fusions (n=481/631; 76.2%; Figure 5A), aneuploid karyotypes (n=68/631; 10.8%), hotspot SNVs (n=34/631; 5.4%) or high confidence gene expression alone (n=48/631; 7.6%). These proportions varied between molecular subtypes (Figure 5A).

**Figure 5:**
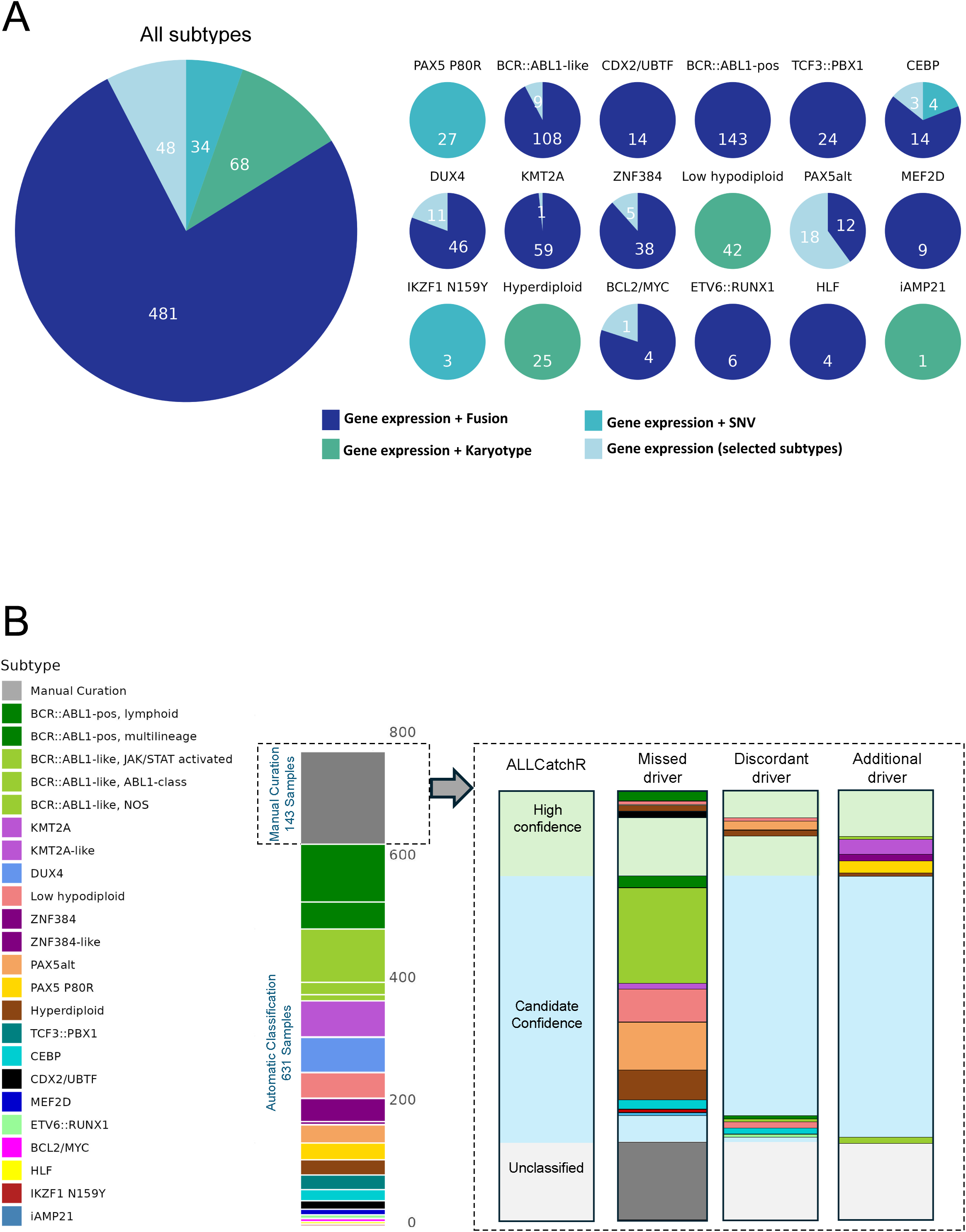
Overview of classification confidence and manual curation. (A) Summary of the basis for classification in cases receiving an unambiguous automatic IntegrateALL subtype assignment for the entire cohort (n=631/774; 81.5%) and individual subtypes. (B) The remaining 143/774 cases (15.5%) were flagged for manual curation. The ALLCatchR confidence categories for gene expression based subtype classification are shown together with the results of manual curation which identified three categories: (i) *Missed driver,* with ‘candidate’ or ‘unclassified’ gene expression-based predictions lacking expected driver events; (ii) *Discordant driver,* representing cases with ‘candidate’ or ‘high confidence’ gene expression-based predictions and driver calls from other subtypes; and (iii) *Additional driver*, where an additional genomic driver from a distinct molecular subtype was identified alongside a subtype definition which would otherwise meet the requirements of unambiguous automatic classification. Validations on three independent cohorts (n=436) are shown in Supplementary Figure S7.

### IntegrateALL identifies novel driver constellations in samples flagged for manual curation

Samples with ambiguous or incomplete matches to the predefined subtype rule set (n=143/774; 18.5%) were flagged for manual curation (Figure 5B). Of these, 28 (19.6%) and 89 (62.2%) had high-confidence or candidate-confidence ALLCatchR subtype allocations, respectively; 26 (18.2%) were unclassified by ALLCatchR and remained so after review. Most flagged samples (n=88/143; 61.5%) lacked a confirming genomic driver call despite an ALLCatchR prediction. This mainly involved subtypes whose original definitions allowed expression-only cases (i.e. *BCR::ABL1*-like (n=32), *PAX5*alt (n=16)), aneuploid subtypes where KaryALL did not recover a matching karyotype (i.e. hyperdiploid (n=12), Low Hypodiploid (n=12)), or other molecular subtypes (n=16). Here, IntegrateALL identifies cases which would either benefit from repeat sampling with higher blast infiltration and/or orthogonal genomic validation.

Discordance between gene expression subtype and driver calls occurred in 13/143 (9.1%) cases. These included samples with candidate or high confidence gene expression subtypes in which drivers from a different subtype were present while the expected driver was absent (n=7; e.g. *PAX5* P80R mutations in cases with PAX5alt, CEBP or *BCR::ABL1* gene expression signatures). One *BCR::ABL1*-like gene expression case without driver fusion was resolved as *BCR::ABL1*-positive after identification of the gene fusion. KaryALL corrected two ALLCatchR hyperdiploid predictions by identifying Near Haploid karyotypes, illustrating its value in separation of these subtypes which share similar gene expression signatures. Conversely, three low hypodiploid cases by expression and manual karyotype review were misclassified as hyperdiploid or iAMP21 by KaryALL.

Importantly, in the remaining 16/143 (11.2%) cases (2.6% of the entire cohort), IntegrateALL identified secondary genomic drivers in addition to the subtype-defining alteration (Figure 6A-B, Supplementary Table S5). Examples included hyperdiploidy co-occurring with KMT2A fusions and *PAX5* P80R or *BCR::ABL1*-like gene fusions in ZNF384, *PAX5* P80R and hyperdiploid cases. Two *BCR::ABL1*-like cases harbored driver fusions from different signaling trajectories (Figure 6B).

**Figure 6:**
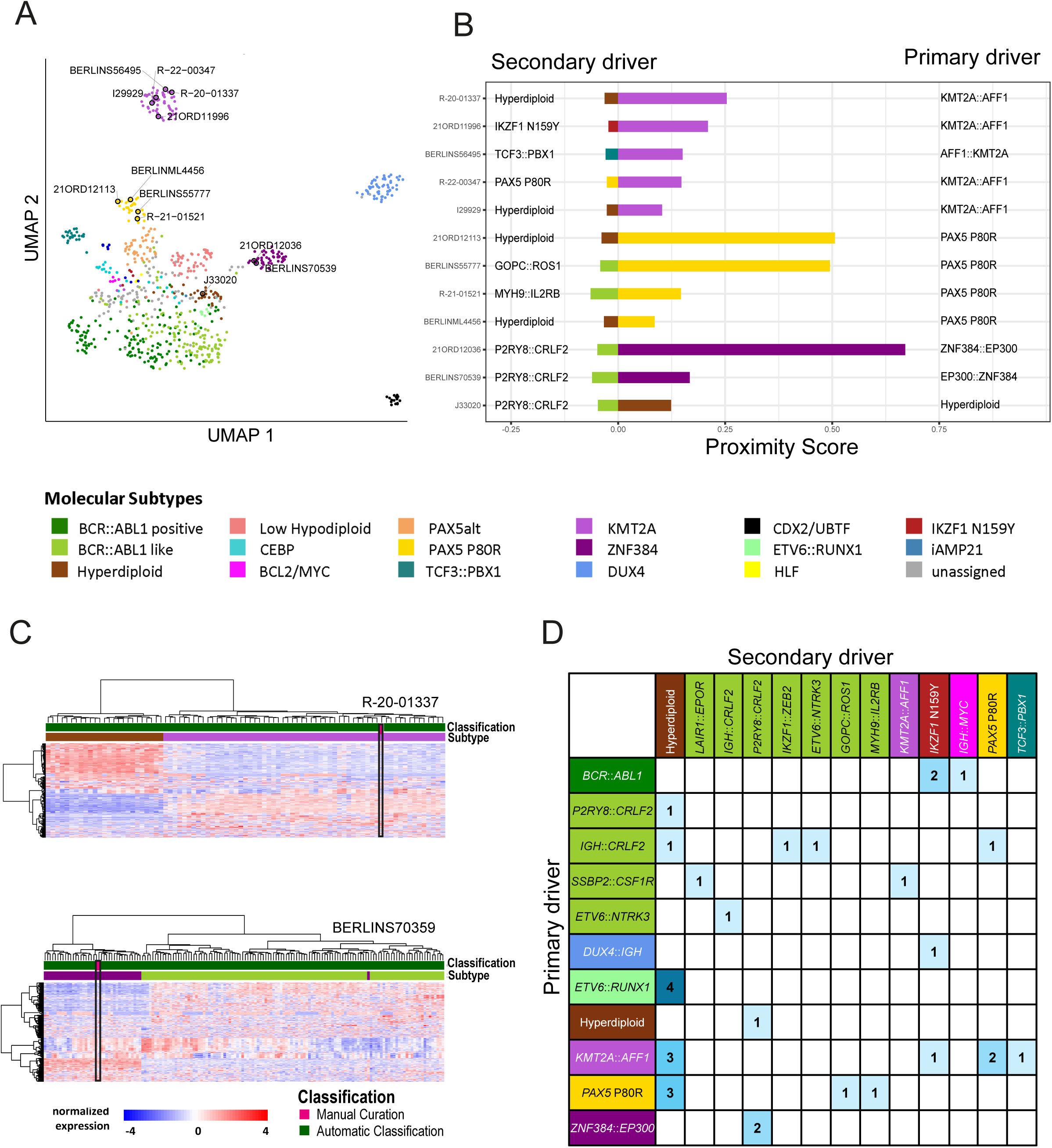
Characterization of double driver cases. (A) UMAP projection of 774 samples based on 2,802 LASSO-selected subtype-specific genes, highlighting double driver cases (n=12) by sample identifiers. (B) Gene expression-based proximity analysis of double driver cases to both molecular subtypes were performed analyzing each double driver case together with all other samples of the two involved subtypes. Proximity scores were calculated using Euclidean distances of all samples of both involved subtypes to the sample of interest using principal component analysis of 600 top variable expressed genes in this sample set. Cases with *BCR*::*ABL1*-like driver fusions from distinct signaling trajectories (e.g., JAK/STAT and ABL-class) were excluded from this analysis. (C) Heatmaps indicate unsupervised clustering (Ward.D2) of gene expression of two representative double driver cases compared to other samples from both involved subtypes. (D) Table illustrating the distribution of genomic drivers identified in double driver cases of the entire B-ALL dataset (n=31/1210; 2.6%). Molecular driver subtypes are characterized as primary or secondary according to the predominating gene expression-based subtype classification.

Notably, ALLCatchR predicted a dominant subtype in nearly all dual-driver samples. UMAP analysis based on 2,802 LASSO-selected, subtype-defining genes^8^ grouped these cases together with one of the ALLCatchR subtypes (Figure 6A). To assess how closely individual double driver samples align with their annotated subtypes, we used the 600 top variable expressed genes from all samples of the two involved subtypes for a subtype-centered principal component analysis (PCA) which compared the position of the double driver sample in PCA space compared to the centroid (mean PC1/PC2 coordinates) of each subtype group. The Euclidean distance to each centroid was calculated to quantify subtype proximity (Figure 6B). This nearest neighbor analysis and unsupervised clustering of sample-level expression profiles (Figure 6C) using the top 600 variable expressed genes, both confirmed dominance of a single driver program.

Blast fraction influences manual curation outcomes. ALLCatchR-inferred blast proportions in discordant (median: 75.2%; p=0.057) or additional-driver cases (median: 77.3%, p=0.09) were comparable to automatically classified samples (median: 73.9%). In contrast, unclassified cases (median: 54.7%; p<0.01) and cases with missed drivers (median: 57.7%; p<0.01) showed significantly lower blast fractions, highlighting reduced sensitivity at low tumor content (Supplementary Figure S6). A similar pattern was observed for virtual karyotyping: samples with expression-based aneuploid subtypes and no KaryALL confirmation had lower predicted blast fractions (median: 60.6%, p>0.001) than automatically classified cases.

For validation, we tested IntegrateALL in 436 independent B-ALL samples from two own cohorts (n=293, including 98 unselected routine-diagnostic cases; Munich Leukemia Laboratory) and one public cohort 233^28^ representing also pediatric patients. In total, 343 (78.7%) cases were automatically classified, and 93 (21.3%) samples were flagged for manual curation, (Supplementary Figure 7), confirming the distribution of our initial test set (Figure 5B). The reasons for manual curation - missed driver (41.9%), discordant driver (14.0%) and additional driver (16.1%) - were comparable, supporting IntegrateALL’s applicability to independent data sets.

### Dual drivers are present in 2.6% of B-ALL and are associated with one dominant gene expression program

To characterize the full spectrum of double driver cases, we analyzed combined data across all cohorts (n=1,210 samples). IntegrateALL identified 31 (2.6%) samples with two driver calls corresponding to two distinct subtype definitions confirming the observations from our first cohort (Figure 6D). Most frequently, hyperdiploidy co-occurred with subtypes, in which it is not commonly described (*ETV6::RUNX1,* n=4; KMT2A, n=3; *PAX5* P80R, n=3; *BCR::ABL1*-like, n=2). Several *BCR::ABL1*-like cases harbored driver fusions from different signaling trajectories (e.g., JAK/STAT and ABL1-class drivers or *NTRK3*; n=4). The *BCR::ABL1*-like driver *P2RY8::CLRF2* occurred in one hyperdiploid case and two *ZNF384* cases. Subtype defining single nucleotide variants were also identified as additional drivers: *IKZF1* N159Y in *BCR::ABL1*-positive (n=2), DUX4 and KMT2A or *PAX5* P80R in KMT2A and *BCR::ABL1*-like ALL, respectively. Consistent with the primary cohort (Figure 6A-B), a single expression program predominated in most dual-driver samples. The distribution of secondary drivers suggests that several aberrations might function as primary drivers but also as cooperating events – which is well established for hyperdiploidy but might also apply to *IKZF1* N159Y, *PAX5* P80R and *P2RY8::CRLF2*.

### IntegrateALL’s interactive report provides user-friendly access to heterogeneous tool outputs

To ensure consistent and intuitive access to the heterogeneous tool outputs, we developed an interactive HTML report (Figure 7). The interface highlights findings relevant to ALL subtype classification while preserving the ability to explore raw results. A navigation bar enables rapid movement between sections. The report includes: a classification summary indicating WHO-HAEM5 / ICC alignment (automated classification) or the need for manual curation; the ALLCatchR gene expression-based subtype allocation; KaryALL classifications for aneuploid subtypes; RNA-Seq quality control metrics; hotspot single nucleotide variant calls; RNASeqCNV-visualizations (including a karyogram summary); and gene fusion calls from ARRIBA and FusionCatcher presented in sortable tables with accompanying visualizations.

**Figure 7:**
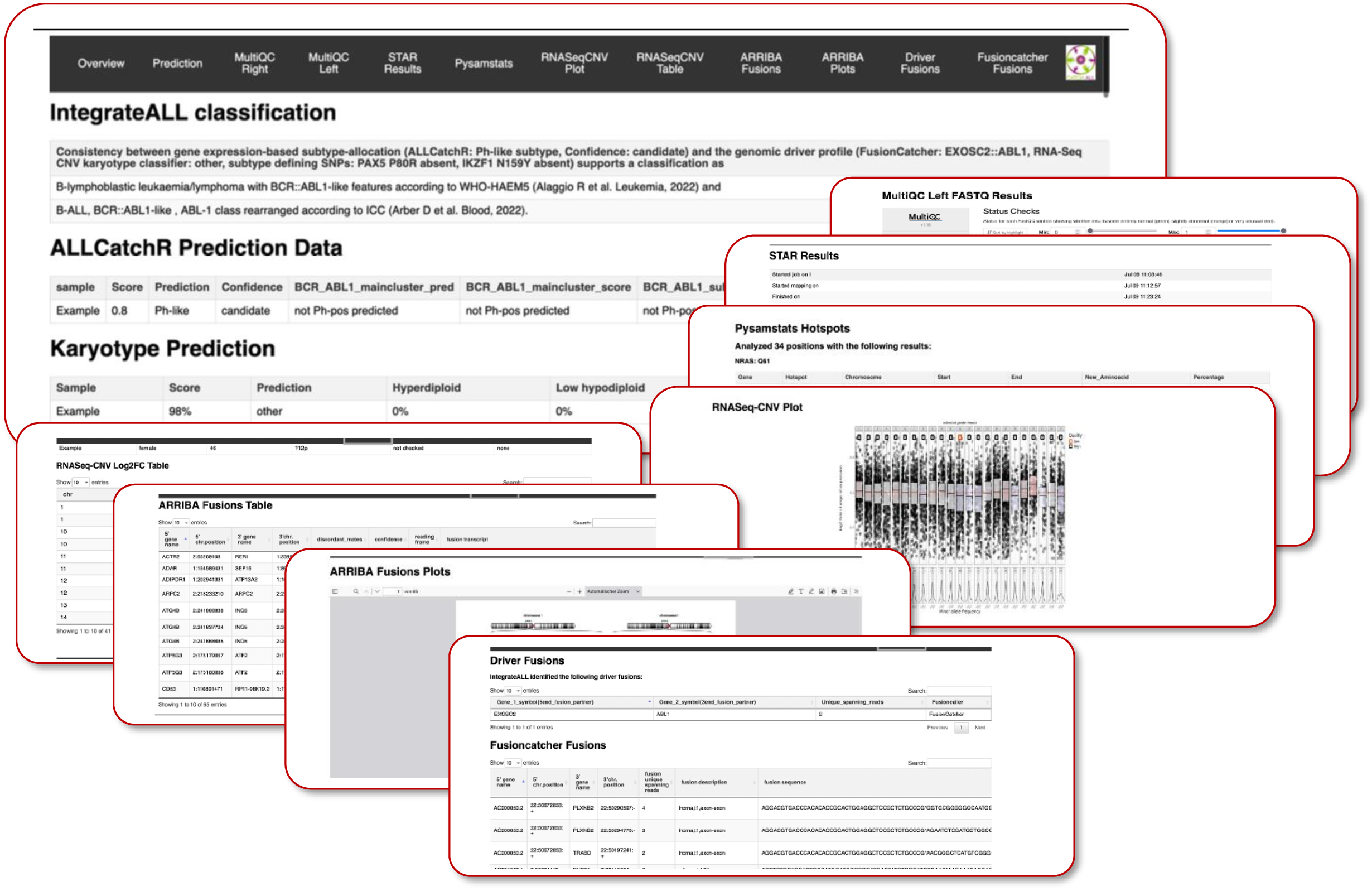
Overview of the IntegrateALL interactive report. To facilitate direct and user-friendly access to all data layers, IntegrateALL provides an interactive HTML document shown here for one example case. The top panel shows the navigation bar, linking to key sections: overview, prediction summary and subtype classification according to ICC and WHO-HAEM5, quality control (MULTIQC), alignment (STAR), SNV analysis (GATK workflow, Pysamstats), copy number profiling (RNASeqCNV; KaryALL) and gene fusion calling (ARRIBA and FusionCatcher). Below, is shown for a concordant case, alongside ALLCatchR and KaryALL predictions. The remaining thumbnails summarize individual analysis modules, providing a visual impression of the integrated output layout.

## Discussion

Recent updates in B-ALL classification have expanded the number of recognized molecular subtypes to a total of 12 according to WHO-HAEM5 and 27 according to ICC, including 5 provisional entities^1,2^. Most of these subtypes are defined by distinct gene expression profiles coupled with specific genomic driver aberrations. Conventional diagnostic approaches require integration of multiple modalities-cytogenetics, FISH, breakpoint PCRs, targeted sequencing - to capture all relevant alterations. This process is time-consuming, costly and may miss cryptic lesions (e.g., *DUX4* fusions) that only manifest as expression signatures. In this context, RNA-seq has emerged a single-platform alternative capable of detecting gene fusions, sequence mutations, virtual karyotypes and the overall gene expression signature in one assay. Various recent studies demonstrate the feasibility and clinical value of RNA-seq–based classification, indicating an improved subtype classification beyond conventional diagnostics^27,30,34,35^.

We and others have developed machine learning classifiers for B-ALL subtype classification (e.g., ALLSorts^9^, Allium^11^ or ALLCatchR^8^). These tools achieve overall classification accuracies of 90-95% and even more in well represented subtypes with clear cut gene expression signatures. However, some rare subtypes remain a challenge, including iAMP21 or near haploid ALL and samples with low blast proportions. Ambiguous cases – especially when harboring multiple drivers – might escape expression-based classification. Frameworks have been established which integrate outputs from gene expression classifiers with second data layers to improve classification with either sentinel driver aberrations form RNA variant calling (MD-ALL^36^, RaScALL^37^ or conventional diagnostics (clinALL^13^) as well as DNA-methylation profiling (ALLIUM^11^). These data underscore the improvements achieved for subtype classification by combining gene expression analysis with underlying drivers or independent functional signatures.

Our newly developed IntegrateALL pipeline fits in this evolving landscape as an open source, end-to-end pipeline rather than a single prediction algorithm. We aim to provide a workflow environment that leverages state-of-the-art methods for unsupervised, systematic identification of molecular drivers, including gene fusions, expression-based subtypes, SNVs, and karyotype abnormalities. For a systematic subtype allocation, we used published descriptions of individual molecular B-ALL subtypes^18,19^ together with comprehensive cohort analyses^16,20,27^ and diagnostic classifications^1,2^ to provide a comprehensive catalogue of genomic B-ALL drivers and corresponding gene expression signatures. An open design allows for adaptation and importantly expansion, as novel B-ALL subtypes are still identified^38^. We do not aim to outperform other existing classifiers, but to deliver a reproducible and transparent workflow that captures the correct subtype in the majority of samples automatically, while flagging atypical cases for manual curation. Validation of IntegrateALL in external cohorts achieved an unambiguous automatic classification in 83.7%, 84.8% or 73.8% of samples, limiting the number of samples for detailed manual review. That approach acknowledges that no single classifier will meet diagnostic standards at handling the molecular complexity of B-ALL. Instead, by requiring concordance between gene expression subtype and corresponding driver lesion for an automatic call, Integrate ALL prioritizes precision and interpretability.

Classification of karyotypes based on gene expression has relied so far on expert curation. Although clear enrichment patterns of non-random chromosomal gains and losses have been described^39,40^, individual cases regularly present deviations from the overall average and large-scale chromosomal gains/losses impart relatively subtle transcriptional effects. To provide a systematic and quantitative analysis of aneuploid karyotypes and chr21 amplification / iAMP21 we established KaryALL, an ensemble machine learning classifier for systematic inference for virtual karyotypes from RNA-Seq data based on RNASeqCNV^14^ outputs. By training on 395 cases with SNParray based karyotypes, KaryALL achieved high cross-validation accuracy (∼98%) and F1-score (∼0.96%) in identifying near haploid, Low Hypodiploid, hyperdiploid and iAMP21 cases. Importantly, KaryALL was able to resolve the ambiguity between Near Haploid and Hypodiploid ALL, which are difficult to separate by gene expression alone. To our knowledge, KaryALL is the first classifier for virtual B-ALL karyotypes providing quantitative confidence in the karyotype call, again aiding in the distinction between clear cut cases and rare candidates which would need orthogonal validations for a correct subtype assignment.

Typically, B-ALL samples harbor a single genomic driver aberration or a subtype-defining aneuploidy pattern. Targeted approaches for B-ALL classification therefore might stop after identifying the first – probably more frequent or more prominent – driver. IntegrateALL flags these cases for manual curation. Application to large cohorts (n=1,210 cases) provides first estimates that 2.6% of B-ALL cases harbor multiple subtype-defining aberrations simultaneously. Co-occurrence of hyperdiploid karyotypes and *ETV6::RUNX1*, *PAX5* P80R, *KMT2A*- or *CRLF2*-rearrangements are not unexpected since hyperdiploidy is also observed in *BCR::ABL1*-positive ALL^33^. However, it is a novel observation that hyperdiploidy can be acquired independently of the underlying driver fusion. Less expected was the co-occurrence of *BCR::ABL1*-like ALL driver fusions (e.g. *CRLF2*, *ROS1* or *IL2RB*) or *EP300::ZNF384* with *PAX5* P80R mutations. *PAX5* P80R has so far been described as an exclusive driver with a subtype defining gene expression signature^17,19,41^. Our data suggests that this mutation might also cooperate with other drivers during leukemogenesis or that multiclonal driver events co-occur in the same patient. Similarly, the subtype defining^42,43^ *IKZF1* N159Y mutation was also observed to co-occur with *BCR::ABL1*, *IGH::DUX4* or *KMT2A::AFF1* gene fusions. IntegrateALL also identified two driver fusions form different signaling types (JAK/STAT-driven plus ABL-class or other) in *BCR::ABL1*-positive / -like cases confirming case reports with similar observations^44–47^. Notably, the *BCR::ABL1*-like driver *P2RY8::CRLF2* seems to be especially prone to co-occurrence with other drivers ^47^.

While these reports are mostly anecdotal, IntegrateALL identifies such double driver cases systematically. Remarkably, our gene expression analyses identified one dominant subtype-specific gene expression signature in most double-driver cases, suggesting that one driver remains in control of oncogenic signaling. Although sample swapping remains a possibility in some of these cases, the non-random enrichment patterns of specific dual drivers seem to be more in line with co-occurrence in B-ALL. In all literature reported cases, patients with double drivers tended to be high-risk and often required combination therapy (e.g. TKI plus chemotherapy)^44,45^, underscoring the need for systematic unsupervised identification.

Research requires reproducible workflows. IntegrateALL provides a systematic pipeline of standard tools used for B-ALL research. The Snakemake environment facilitates implementation of novel applications for specific research questions which can then build upon a systematic and comparable baseline characterization. Free accessibility of all raw outputs enables unsupervised analysis beyond established subtype definitions and drivers. An open design allows for seamless implementation of novel classification rules.

In conclusion, the integration of machine learning into the analysis of B-ALL RNA-Seq data represents an important step toward a more systematic and data-driven classification of leukemia subtypes. IntegrateALL facilitates diagnostic workflows by enabling large-scale, standardized profiling while simultaneously identifying complex cases that require manual curation. By incorporating gene expression-based subtype classification, karyotype prediction, and secondary driver detection, the pipeline provides a comprehensive framework for refining the molecular characterization of B-ALL.

The results demonstrate that computational models can enhance the accuracy and consistency of subtype classification while also capturing additional genomic complexity that may influence disease progression. The ability to distinguish primary from secondary drivers further supports a more nuanced interpretation of cases with multiple alterations. These findings highlight the potential of machine learning-assisted approaches to improve leukemia diagnostics and provide a scalable method for integrating diverse genomic data in clinical and research settings.

## Data and Code availability

Published data used in the manuscript can be accessed through^16,17^. Newly generated RNA-Seq and SNParray data will be provided upon acceptance of the manuscript in a peer-reviewed journal.

The code used in this work is available in a publicly accessible repository at https://github.com/NadineWolgast/IntegrateALL under the MIT License, ensuring open access for modification and distribution. For detailed guidance on implementing and running the pipeline, users can refer to the comprehensive instructions provided in the repository’s README file, available at https://github.com/NadineWolgast/IntegrateALL/tree/main#readme.

## Acknowledgments

The large language model ChatGPT (OpenAI, USA) was used for language editing without interfering with any content of the manuscript. The large language model Claude (Anthropic, USA) was used for code refactoring and optimization without interfering with any scientific content or methodology.

## Author contribution

NW, TB, AMH, CDB, and LBas designed the study. NW and LBas established the ruleset for molecular subtype classification according to WHO-HAEM5 and ICC. NW, TB and LBas developed the karyotype classifier. NW and LBas validated the ground truth of RNASeqCNV using SNParray data. BTH, KI and MJB provided data and analyses for establishing the genomic ground truth and performed wetlab validation. NW and MM established the Snakemake pipeline framework, NW integrated and validated the pipeline tools. TB, MM, AMH, JK, SB, WW and SH validated the pipeline for distribution. WW, CH, SW, AC and MB contributed B-ALL sequencing data and validated ground truth and/or contributed to the classifier concept. MN, NG and MB contributed to the classifier concept and its real-world applicability. LBas, AMH and CDB supervised the project. NW, AMH, CDB, and LBas drafted the first version of the article. All authors revised and approved the final version of the article.

