## Supplementary Figures for "IntegrateALL: an end-to-end RNA-seq analysis pipeline for multilevel data extraction and interpretable subtype classification in B-precursor ALL"

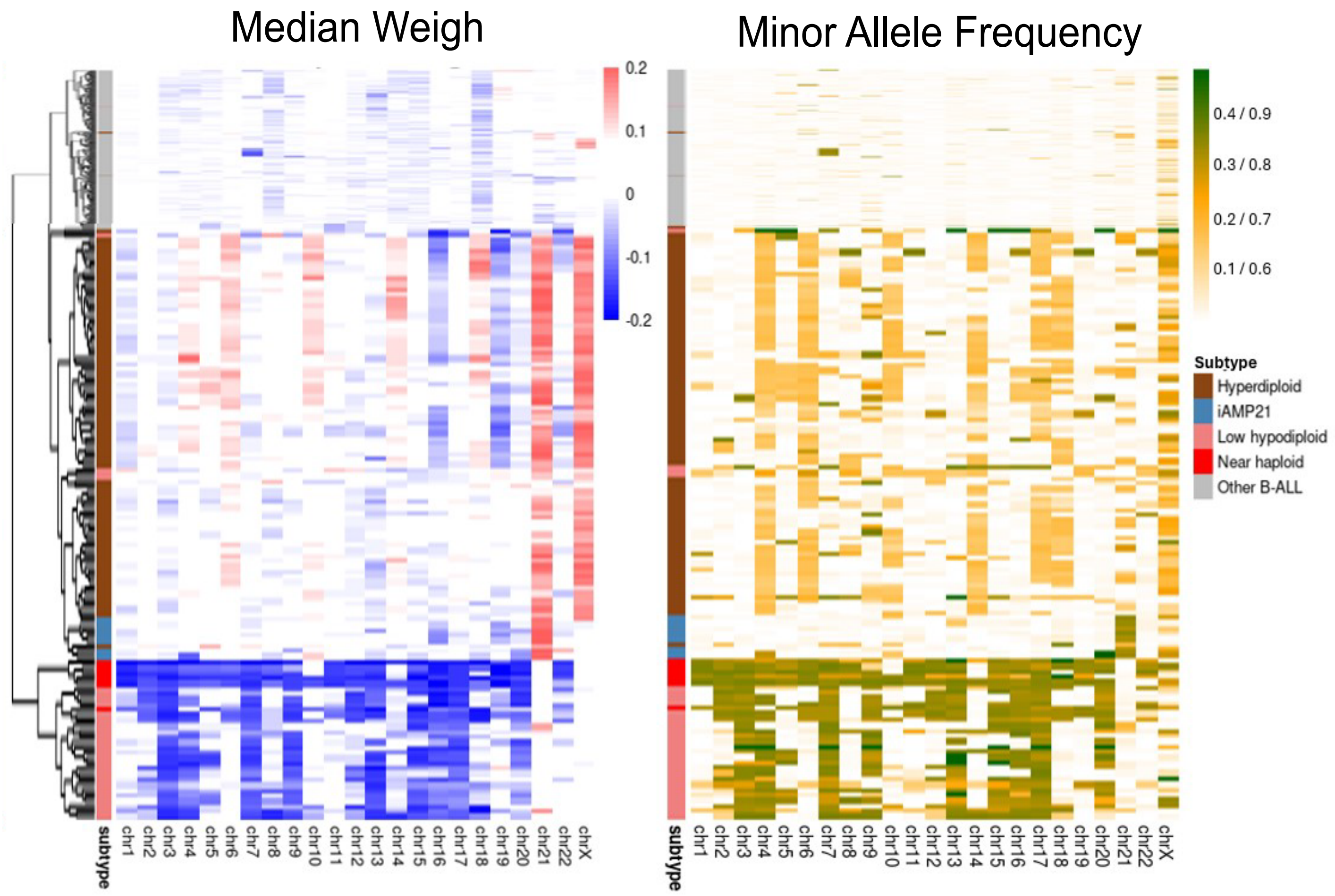


**Supplementary Figure 1:** Heatmaps of RNASeqCNV-derived data (median weight and minor allele frequency) across the training cohort (n=395), annotated by subtype. Gains appear in red/yellow; losses in blue/green. Clustering using Ward.D2 on expression data reveals distinct subtype separation; frequent subtypes were compressed.

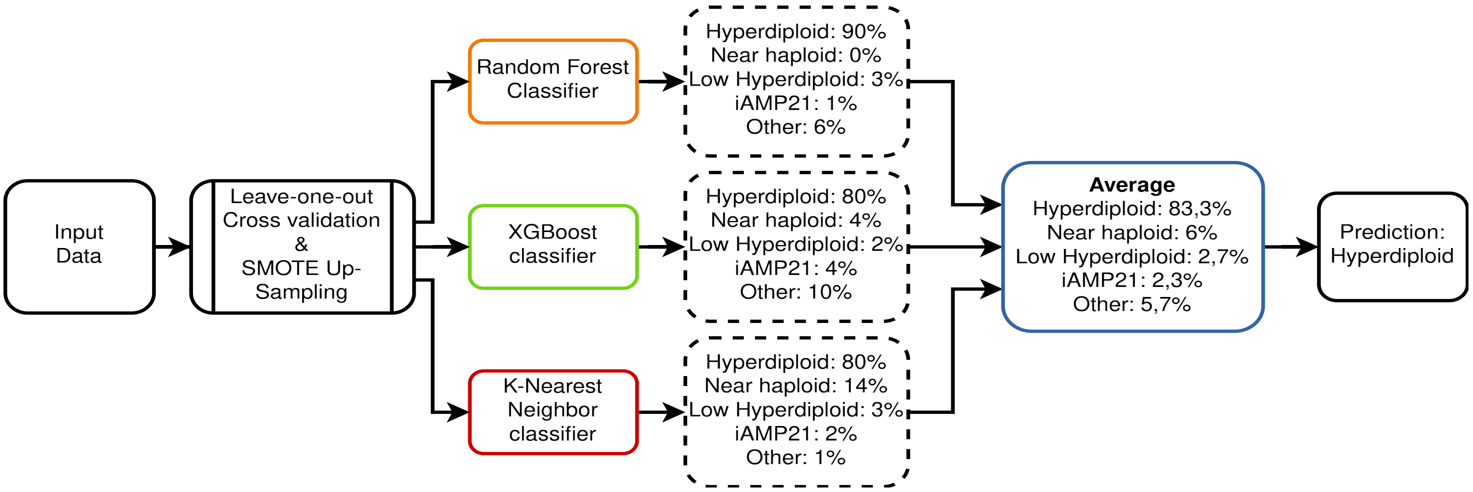


**Supplementary Figure 2:** Workflow of the KaryALL classifier. Input features from RNASeqCNV are processed using SMOTE up-sampling and leave-one-out cross-validation with a voting ensemble (Random Forest, XGBoost, k-NN).

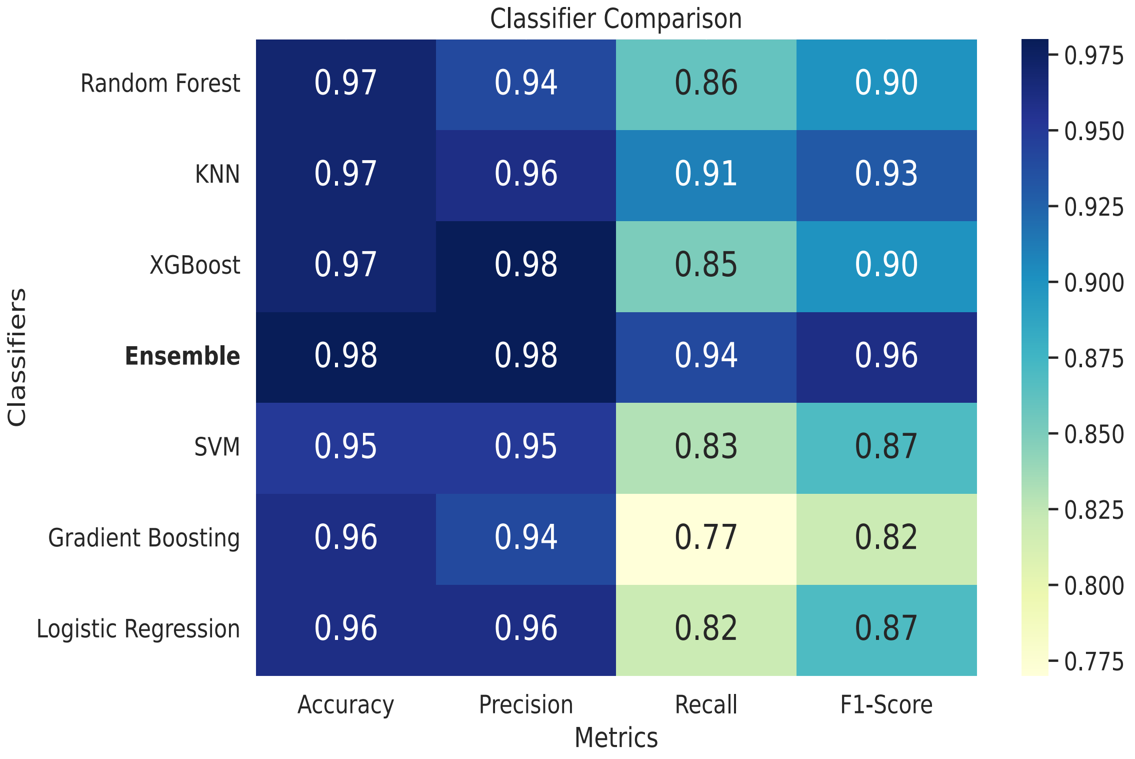


**Supplementary Figure 3:** Classifier comparison. Performance comparison of six single classifiers and the ensemble model using accuracy, precision, recall, and F1-score; the ensemble outperforms all individual models.

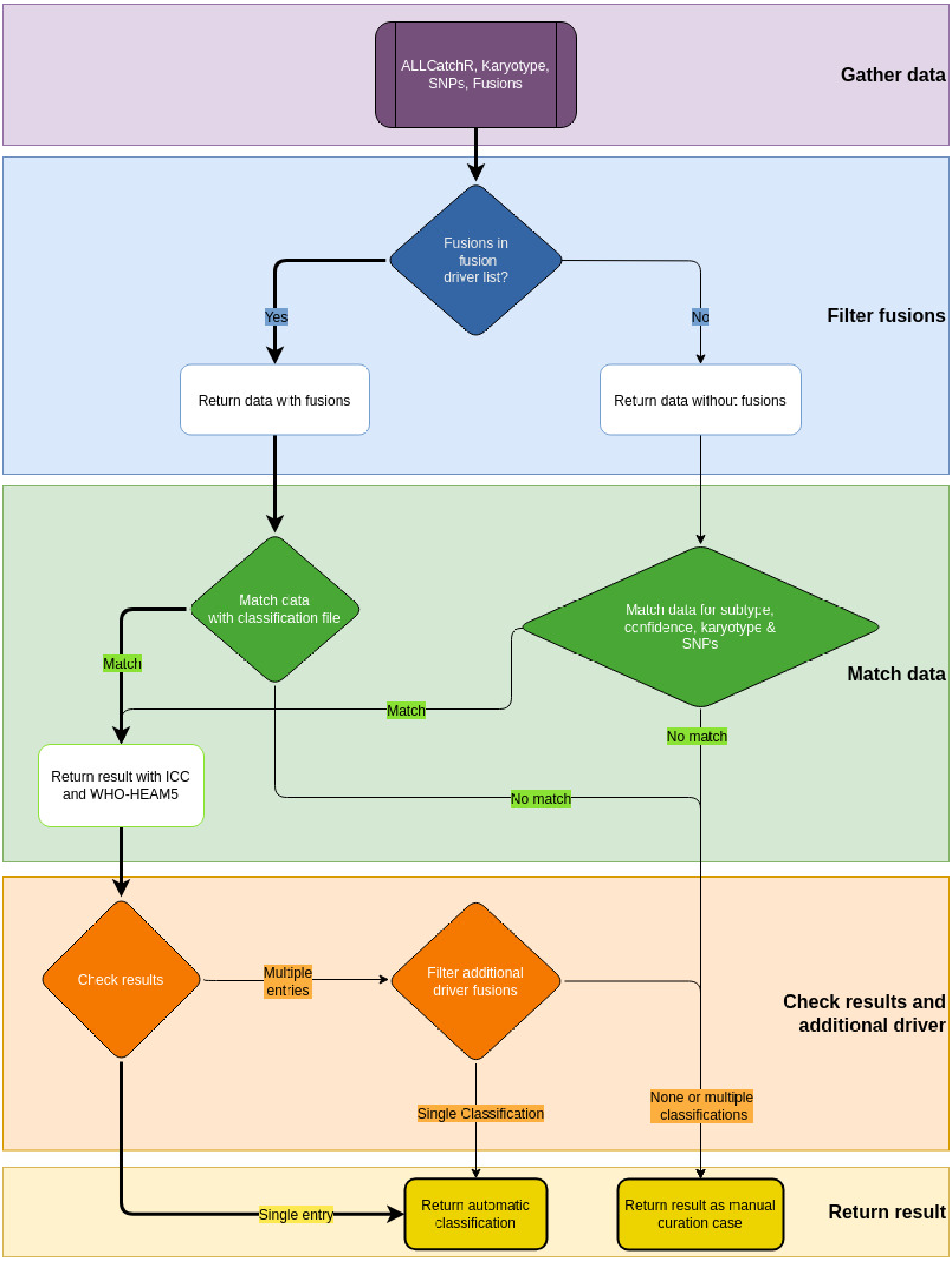


**Supplementary Figure 4:** Schematic overview of the IntegrateALL classification algorithm to assign molecular subtypes based on multi-layered genomic and transcriptomic data. The flowchart is organized into five color-coded levels, labeled on the right margin: Gather data (purple), Filter fusions (blue), Match data (green), Check results and additional drivers (orange), and Return result (yellow). Data inputs include fusion calls, gene expression profiles, SNV hotspots, and RNASeqCNV-based virtual karyotypes. Fusion results are first filtered against a curated list of subtype-defining drivers. Fusion-positive samples are matched to a classification reference to assign an ICC and WHO-HAEM5 subtype; unmatched cases proceed to further evaluation. Fusion-negative samples are assessed for concordance between expression-based subtype, prediction confidence, karyotype, and SNVs. If predefined criteria are met, diagnostic labels are assigned; otherwise, the case is flagged for manual curation. In the decision phase preceding classification, the algorithm determines whether a unique classification can be made. If multiple potential classifications or additional drivers are detected, further filtering is applied. The final decision returns either an automatic classification or flags the case for manual review.

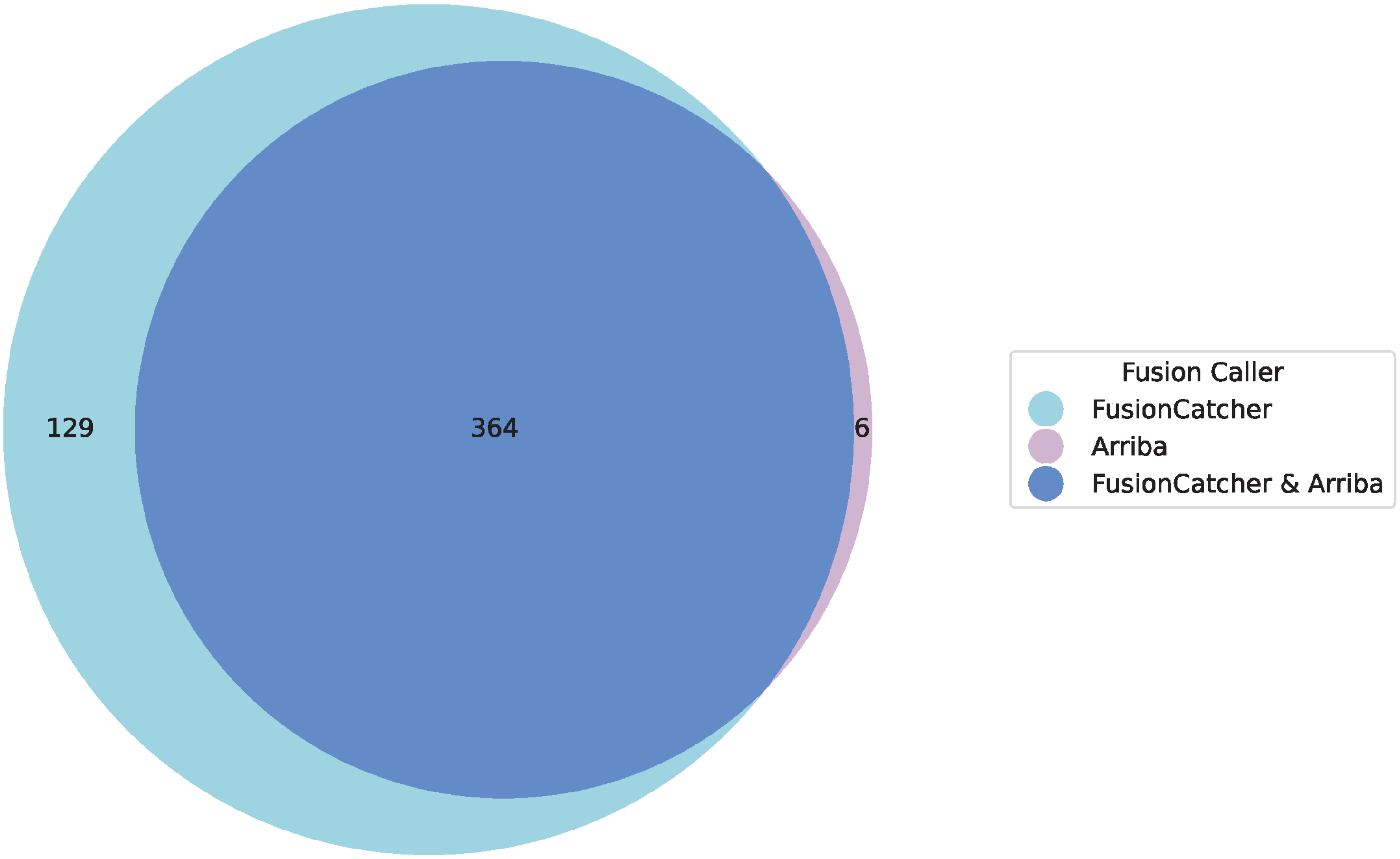


**Supplementary Figure 5:** Fusions called by FusionCatcher and Arriba. Supplementary Figure 5 shows the overlap of gene fusion calls obtained using FusionCatcher and Arriba across the cohort of 774 samples.

In total, 364 fusions were identified by both algorithms, while 129 were detected exclusively by FusionCatcher and 6 only by Arriba. Notably, certain fusions such as *ATXN7L3*::*UBTF* and *IGH*::*DUX4* were identified only by FusionCatcher.

The full list of fusions is provided in Supplementary Table S4.

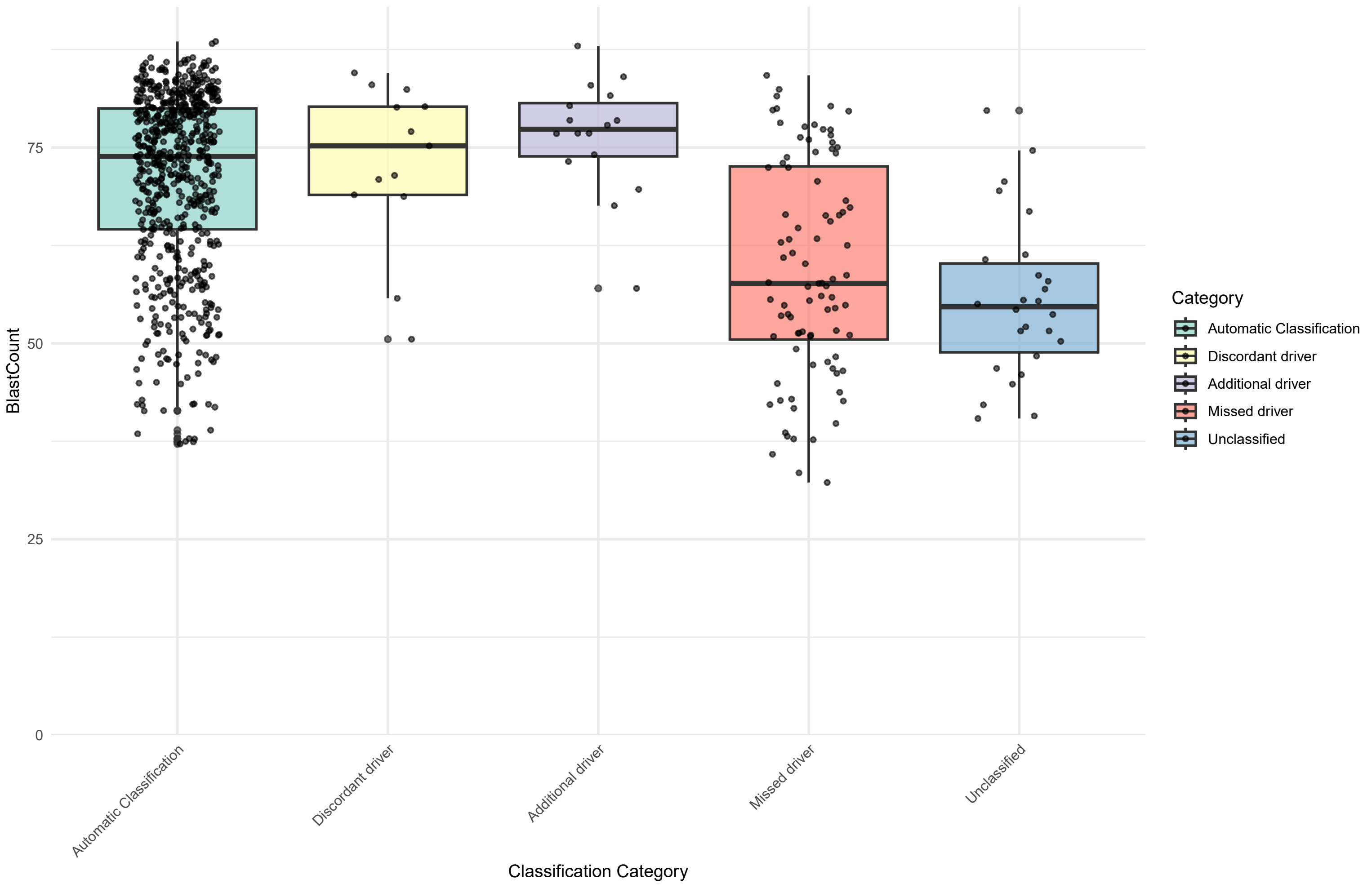


**Supplementary Figure 6:** Blast Count comparison of automatic classified samples vs. manual curation samples for each category.

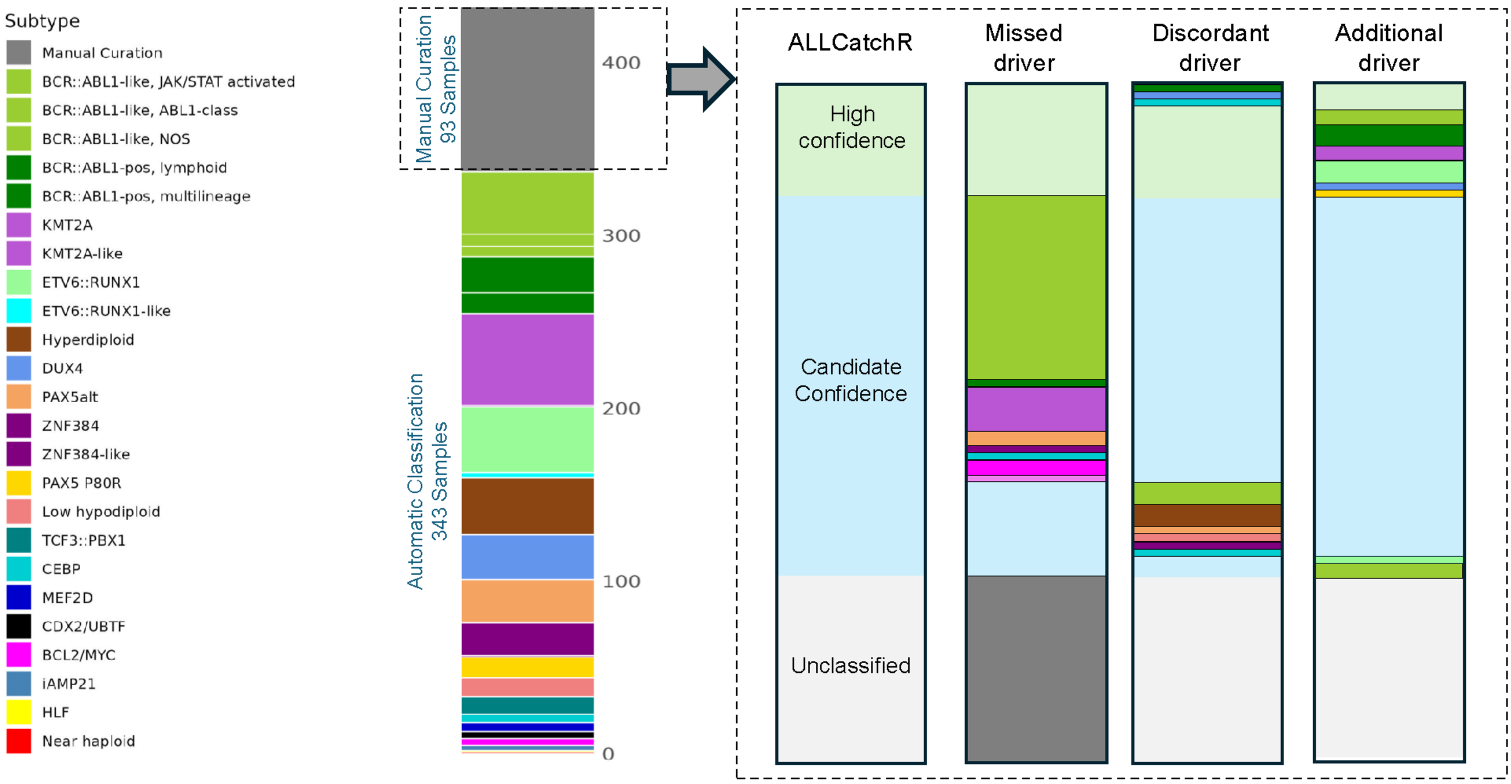


**Supplementary Figure 7:** External cohort validation of 95/436 manually flagged cases shows similar classification patterns to our internal cohort (Figure 5B).
